# Highly sensitive *in vivo* detection of dynamic changes in enkephalins following acute stress

**DOI:** 10.1101/2023.02.15.528745

**Authors:** Marwa O. Mikati, Petra Erdmann-Gilmore, Rose Connors, Sineadh M. Conway, Jim Malone, Justin Woods, Robert W. Sprung, R. Reid Townsend, Ream Al-Hasani

## Abstract

Enkephalins are opioid peptides that modulate analgesia, reward, and stress. *In vivo* detection of enkephalins remains difficult due to transient and low endogenous concentrations and inherent sequence similarity. To begin to address this we previously developed a system combining in vivo optogenetics with microdialysis and a highly sensitive mass spectrometry-based assay to measure opioid peptide release in freely moving rodents (Al-Hasani, 2018, eLife). Here we show improved detection resolution and stabilization of enkephalin detection, which allowed us to investigate enkephalin release during acute stress. We present an analytical method for real-time, simultaneous detection of Met- and Leu-Enkephalin (Met-Enk & Leu-Enk) in the mouse Nucleus Accumbens shell (NAcSh) after acute stress. We confirm that acute stress activates enkephalinergic neurons in the NAcSh using fiber photometry and that this leads to the release of Met- and Leu-Enk. We also demonstrate the dynamics of Met- and Leu-Enk release as well as how they correlate to one another in the ventral NAc shell, which was previously difficult due to the use of approaches that relied on mRNA transcript levels rather than post-translational products. This approach increases spatiotemporal resolution, optimizes the detection of Met-Enkephalin through methionine oxidation, and provides novel insight into the relationship between Met- and Leu-Enkephalin following stress.

## Introduction

Endogenous opioid peptides are released at significantly lower concentrations than small molecule neurotransmitters and undergo post-translational processing that yields ∼30 unique peptides from 4 precursor molecules. Together, these properties have made sensitive peptide detection difficult to achieve with existing technologies [1]. Here, we describe an analytical method that enables quantification of dynamic *in vivo* changes in both Leu- and Met-enkephalin (Leu-Enk and Met-Enk) peptides in rodents. We show unique release profiles between Leu- and Met-Enk in response to two different stressors and gain new insight into peptide release dynamics; additionally, our method offers improved spatiotemporal resolution. More broadly, this approach will enable an understanding of how neuropeptide levels change in response to acute or chronic behavioral, pharmacological, and neurophysiological manipulations.

Enkephalins can act on both the delta and/or mu opioid receptors, but their activity often depends on receptor and peptide expression patterns, which differ across the brain [2–5]. Confirming an opioid peptide’s identity based on the activity of the receptor alone poses difficulties. For example, in a single brain region, the mu opioid receptor may be activated by an enkephalin or endorphin peptide [3]. Therefore, any technique that relies on a readout of mu opioid receptor activity is not sufficient to determine whether the peptide is an endorphin or an enkephalin. It has been particularly challenging to measure Met- and Leu-Enk selectively and independently *in vivo*; in fact, both peptides are typically referred to as ‘enkephalins’, which oversimplifies their distinct properties. Though antibody-based techniques [6–9] or liquid chromatography/mass spectrometry (LC-MS) [10–12] have improved Met- and Leu-Enk detection somewhat, limitations such as spatiotemporal restrictions and poor selectivity between enkephalins remain. Our group previously developed real-time *in vivo* detection of evoked opioid peptide release using microdialysis/nanoLC-MS (nLC-MS) after photostimulation [13]. From this strong foundation, we have now optimized a novel opioid peptide detection technique. For the first time, we can selectively measure real-time *in vivo* levels of basal (non-evoked) peptide release, as well as during behavioral manipulations. Our primary goals in establishing this analytical workflow to measure opioid peptide dynamics *in vivo* were to 1) increase temporal resolution allowing for the detection of dynamic changes to short external stimuli and acute conditions, 2) lower the detection threshold to allow for accurate measurements from small brain regions using less volume of sample, 3) measure endogenous release of peptides in the native state without the need for photo- or chemogenetic stimulation, 4) report raw values of measured concentrations rather than % baseline measurements, 5) enable real-time, simultaneous detection of both Met- and Leu-Enk. To meet these goals and measure both Leu- and Met-Enk we used an improved and specialized microdialysis technique coupled with nLC-MS.

## Results

### Optimizing a method for the simultaneous *in vivo* detection of Met- and Leu-Enkephalin

Our workflow (**Fig. 1A**) starts with stereotaxic surgery to implant the microdialysis probe in the mouse nucleus accumbens shell (NAcSh). Along with others, we have shown that the opioid receptor system in the NAcSh mediates both reward and aversion through photostimulation of peptide-expressing neurons across the dorsal/ventral axis [14–20]. Our aim was to measure NAcSh peptide release in response to aversive, stressful stimuli. We use custom-made microdialysis probes, intentionally modified so they are similar in size to commonly used fiber photometry probes thus causing comparable levels of tissue damage (**Fig. 1B**). Importantly, the membrane that sits above the brain region of interest is smaller in diameter when compared to a photometry probe (**Fig. 1B**). After recovery, we begin microdialysis collection every 13 minutes, circulating artificial cerebrospinal fluid (aCSF) and collecting interstitial fluid (ISF) at a flow rate of 0.8 μL/min. After each sample collection, we add a consistent known concentration of isotopically labeled internal standard of Met-Enk and Leu-Enk of 40 amol/sample to the collected ISF for the accurate identification and quantification of endogenous peptide. These internal standards have a different mass/charge (m/z) ratio than endogenous Met- and Leu-Enk. Thus, we can identify true endogenous signal for Met-Enk and Leu-Enk (**Suppl Fig. 1A,C**) versus noise, interfering signals, and standard signal (**Suppl. Fig. 1B,D**). Importantly, we have resolved one of the main barriers preventing the accurate quantification of Met-Enk: the variable oxidation of the methionine residue during sample processing, which results in the signal for Met-Enk being split among multiple m/z values thereby impairing sensitivity. In order to ensure that the signal for Met-Enk is represented by a single m/z value, we converted all methionyl residues to the doubly oxidized sulfone prior to nLC-MS analysis, which allows us to reliably quantify Met-Enk and differentiate it from Leu-Enk without compromising the detection of either peptide [21]. Beyond opioid peptides, this approach represents a key advance and can be applied to any peptide containing a methionine residue. As seen in **Fig. 1C**., before the methionine oxidation reaction at time 0, Met-Enk can be observed in unoxidized, and singly- and doubly-oxidized forms with varying intensities. However, after the reaction is complete at time 1, Met-Enk is observed as doubly oxidized with much higher intensity enabling accurate quantification.

**Figure 1:**
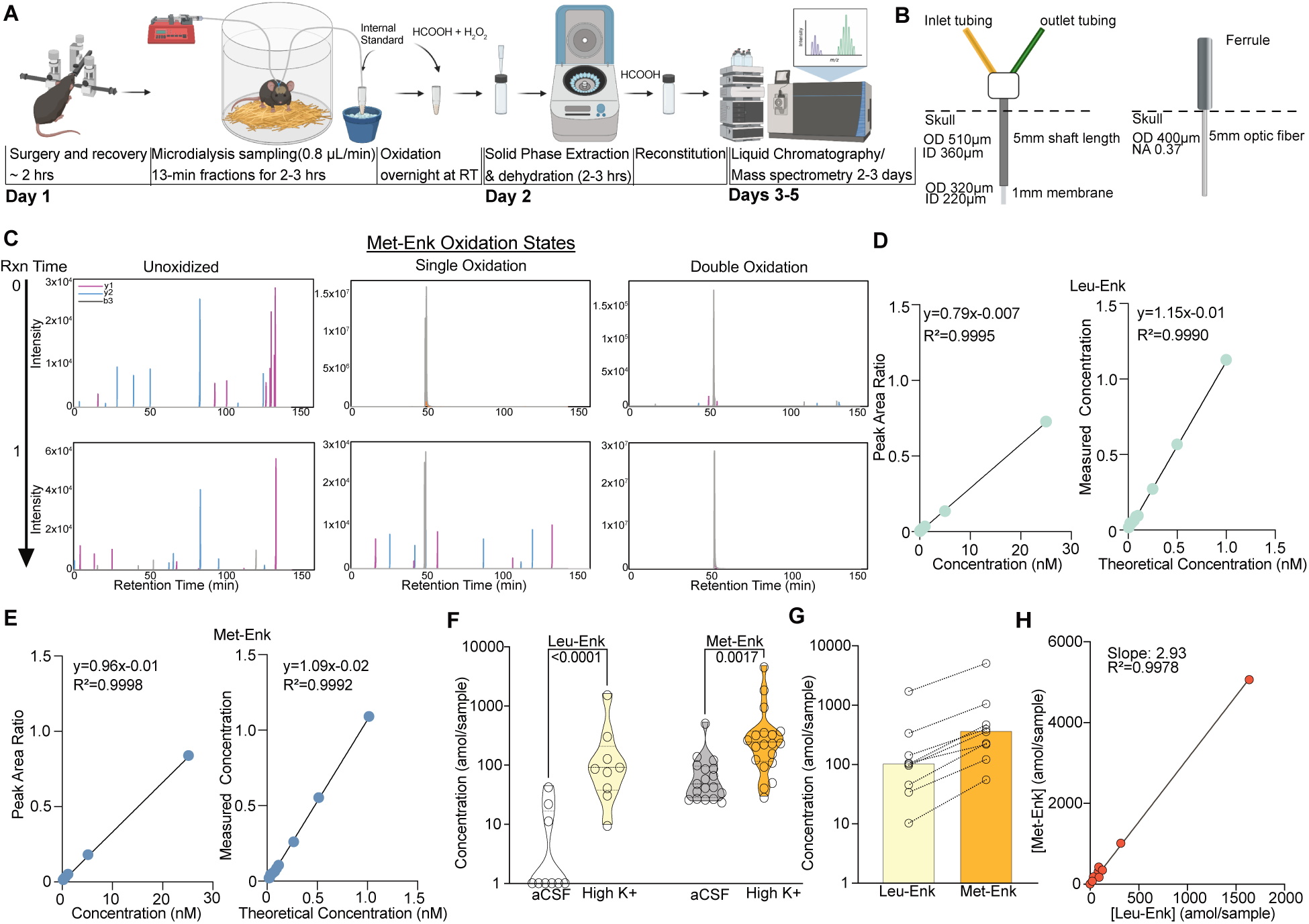
An optimized approach for *in vivo* Met- and Leu-Enk measurement. **A.** Timeline of *in vivo* sample collection on day 1 and methionine oxidation reaction overnight, sample processing on day 2, and data acquisition on the LC-MS, days 3-5. The microdialysis probe is implanted via stereotaxic surgery in the Nucleus Accumbens Shell. The mouse is allowed to recover before being connected to the microdialysis lines. Samples are then collected at a rate of 0.8 μL/min for 13 minutes each. After collection is completed, the samples are oxidized overnight. On day 2 the samples undergo solid phase extraction (SPE) and are then dehydrated and reconstituted using formic acid (HCOOH) before being acquired on the LC-MS. **B**. Custom microdialysis probe specifications including membrane size and inner and outer diameters (ID and OD respectively) compared to fiber photometry probe specifications including OD of optic fiber and numerical aperture (NA). **C**. Before the methionine oxidation at reaction (Rxn) time 0, Met-Enk exists in three different forms with varying intensities, unoxidized (multi-peak), singly oxidized (multi-peak), and doubly oxidized (single peak). After the reaction completes (Rxn time 1), most of the detected signal is in the doubly oxidized form and shows a single peak (>99% signal intensity). Y1, y2, b3 refer to the different elution fragments resulting from Met-Enk during LC-MS. **D.** (Left) Forward calibration curve of Leu-Enk and Met-Enk showing the peak area ratios as the light standard levels are varied. (Right) Reverse calibration curve of Leu-Enk showing the relationship between the heavy standard concentration applied and the measured concentration based on the instrument. **E.** Same setup as **D** but for Met-Enk**. F.** Violin plots showing that high K+ ringer’s solution increases the release of both Leu-Enk and Met-Enk compared to baseline levels in artificial cerebrospinal fluid (aCSF) (Leu-Enk n=9, Met-Enk n=18). The dashed center line indicates the median. **G.** The evoked concentrations of Met-Enk to Leu-Enk in the same samples show that Met-Enk is consistently released at a higher level than Leu-Enk (n=9). **H.** Met-Enk is released at a factor of 2.97 that of Leu-Enk as shown by linear regression analysis of the data in f, suggesting a linear relationship between the two peptides. Data in **F**, **G**, and **H** are transformed to a log scale. In **F**, 2-way ANOVA analysis shows a main effect of peptide (F (1, 51) = 32.30, *p*<.0001), solution (F (1, 51) = 50.04, *p*<.0001), and interaction (F (1, 51) = 9.119, *p=.0039)*. The *p*-values reported were calculated using a Sidak’s multiple comparisons test. Mean difference between baseline Leu-Enk and high K+ 1.533, Mean difference between baseline Met-Enk and high K+ 0.6157, 95% confidence interval 0.3074 to 1.527 (log_10_ values).

To increase the signal to noise ratio, we also performed solid-phase extraction after the oxidation reaction which removes any salt from the collected samples. To determine the lower limit of quantification (LLOQ), forward and reverse curves were created using varying concentrations of the Met- and Leu-Enk standards in accordance with Clinical Proteomics Tumor Analysis Consortium (CPTAC) guidelines [22]. We determined that the lower limit of quantification (LLOQ), for both Leu-Enk and Met-Enk is 40 amol/sample (**Fig. 1D-E**). The improved quantification limit means increased temporal resolution. We now need less volume, (10 μL or less), and shorter sampling times: 10 minutes or less, rather than 15-20 minutes as compared to previous studies [13,23]. To test the utility of the method in measuring real-time changes in peptide levels *in vivo*, we evoked the release of the peptides using high K^+^ ringer’s solution, which depolarizes the cells and causes the release of dense core vesicles containing neuropeptides [24]. This depolarization state significantly increased the release of Met- and Leu-Enk in the NAcSh as compared to baseline **Fig.1F**. Interestingly, Leu-Enk showed a greater fold change compared to baseline than did Met-Enk with the fold changes being 28 and 7 respectively based on the data in **Fig.1F**. Previous studies reporting opioid peptide measurements rarely represent raw concentrations measured. Instead, most past studies reported changes in peptide concentrations as % of baseline [10,13,23]. One key advantage of our improved method is that we can detect reliable measurements of both Met- and Leu-Enk at baseline reported as concentrations in the amol/sample range. We then quantified the ratio of the Met- to Leu-Enk after applying high K+ solution and showed that this ratio is 3:1 in the NAcSh (**Fig. 1G-H**). The release of Met-Enk is postulated both in vitro and ex vivo to be between 3 to 4 times that of Leu-Enk [25–27]. We are the first to not only corroborate the predicted ratio *in vivo* in freely behaving animals, but specifically in the NAcSh. Interestingly, this 3:1 ratio is almost identical to that previously reported in homogenized human tissue from the sympathoadrenal systems using radioimmunoassays [28]. Importantly, this conserved ratio highlights the usefulness of rodent models for investigating the highly conserved opioid system and demonstrates its potential in translation to humans.

### Experimenter handling drives the release of Met- and Leu-Enkephalins in the NAcSh

To highlight the applicability and value of this improved approach, we investigated the changes in the levels of enkephalins following acute stress. It has been hypothesized that enkephalins act as anti-stress signals based on studies that have shown increased preproenkephalin transcript levels following acute stress [29–32]. These studies have included regions such as the locus coeruleus, the ventral medulla, the basolateral nucleus of the amygdala, and the nucleus accumbens core and shell. Studies using global knockout of enkephalins have shown varying responses to chronic stress interventions where male knockout mice showed resistance to chronic mild stress in one study, while another study showed that enkephalin-knockout mice showed delayed termination of corticosteroid release. [33,34]. Moreover, decreased enkephalin expression in the NAc was correlated with an increase in the susceptibility to social defeat [35]. This body of literature supports the hypothesis that enkephalins participate in stress encoding and reactivity. We were specifically interested in how enkephalins mediate acute stress responses. Our group has also recently demonstrated that the photostimulation of a subpopulation of peptidergic neurons in the NAcSh drives aversion [14], which further suggests a role for neurons in the NAcSh during aversive stimuli such as stress. Despite the wealth of data implicating these peptides in the stress response, it has not been possible to measure the dynamics of real-time peptide release in response to acute stressful stimuli. We sought to test the hypothesis that Met- and Leu-Enk are released in the NAcSh following acute stress. Using the nLC-MS detection method, we successfully monitored dynamic changes of Met- and Leu-Enk during two different stressors. The first stressor was defined as experimenter handling, which involves the experimenter scruffing the mouse while connecting to the dialysis set up (**Fig. 1A**). The second stressor was exposure to predator odor (i.e., fox urine).

We show that both Met- and Leu-Enk are released following experimenter handling, and that the levels decrease within minutes after exposure to the stress (**Fig. 2A**). We also analyzed the data with sex as a main effect and did not find any differences between males and females after correcting for multiple comparisons (**suppl. Fig. 2A**). We corroborate our finding from **Fig. 1G-H** showing that the ratio of 3:1 Met-Enk to Leu-Enk is conserved following experimenter handling (**Fig. 2C**). Another key finding is that peptide release appears to be precisely controlled. We show that the amol/sample of Met-Enk released at the beginning of the experiment is predictive of later release. For example, a higher level released (in the fmol/sample range) during the first collections results in lower levels released (in the amol/sample range) during later collections and vice versa (**Fig. 2D**). We show a negative correlation (∼ -0.3) between the first two samples (∼26 minutes) and the later timepoints, samples 3-10 (∼104 minutes). Importantly, this relationship is not due to Met-Enk depletion as the circulation of high K+ aCSF is sufficient to drive the release of Met-Enk at the end of an experiment and reverse the negative correlation (**Fig. 2E**).

**Figure 2:**
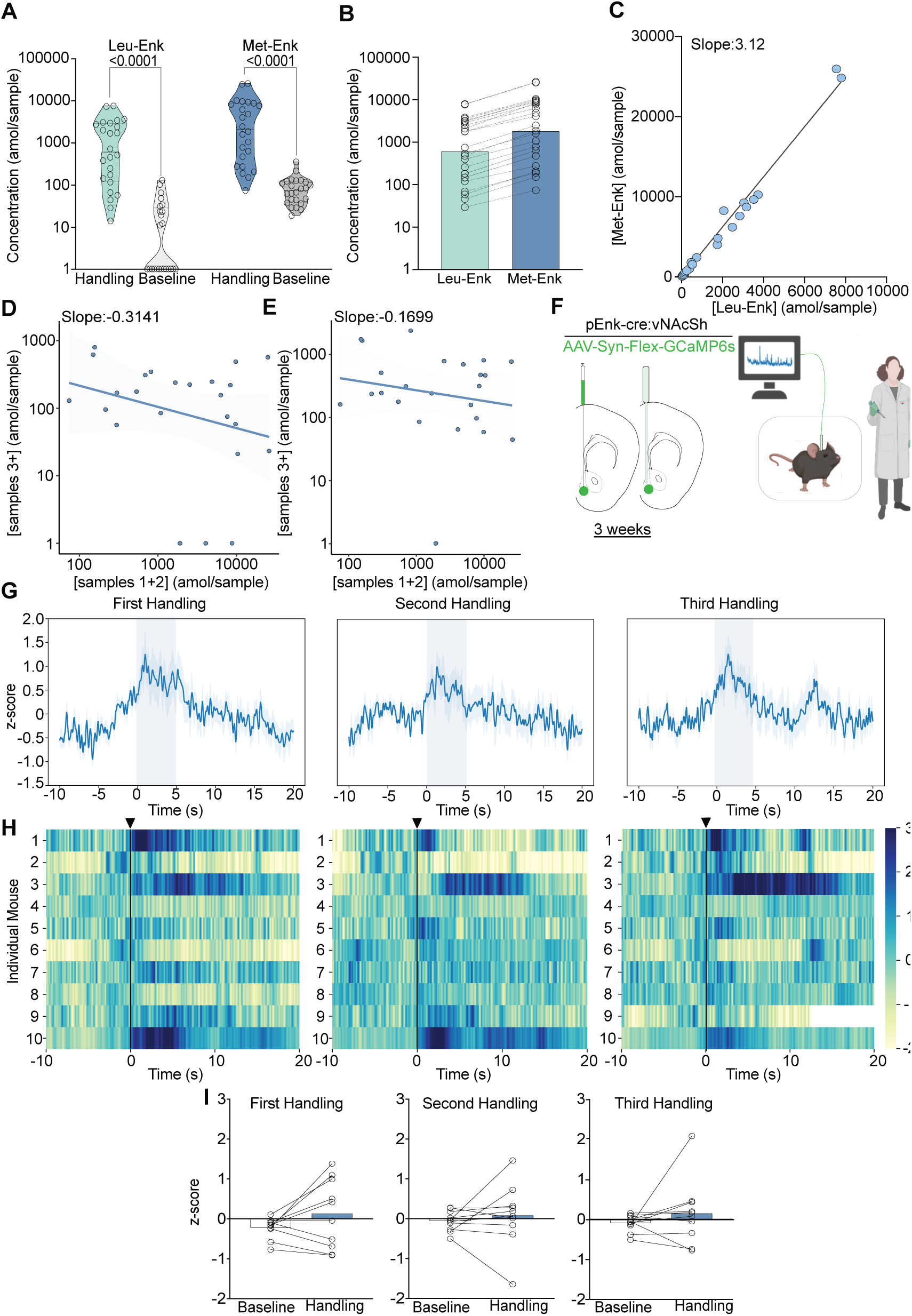
Experimenter handling evokes the release of Met- and Leu-Enk in the NAcSh. **A.** Experimenter handling during microdialysis causes a significant increase in the release of Leu- and Met-Enk in comparison to baseline. 2-way ANOVA on log-transformed data showing a significant effect of peptide F(1,92)=32.58, *p*<0.0001, handling F(1,92)=146.1, *p*<.0001, and interaction F(1,92)=4.778, *p*=.0314, Sidak’s multiple comparisons test was conducted and *p*-values are shown on the figure. Mean difference between baseline Leu-Enk and handling 2.055, Mean difference between baseline Met-Enk and handling 1.426, 95% confidence interval 0.05751 to 1.201 (log_10_ values).**B.** During experimenter handling, Met-Enk is consistently released at a higher rate than Leu-Enk in the same samples. **C.** Linear regression analysis of the data in **B** shows that Met-enk is released at a rate of 3.12 times the rate of Leu-Enk during experimenter handling suggesting a linear relationship between the two peptides. **D.** A negative correlation (-0.3141) shows that if a high concentration of Met-Enk is released in the first two samples, the concentration released in later samples is affected; such influence suggests that there is regulation of the maximum amount of peptide to be released in NAcSh. **E.** The negative correlation in panel d is reversed by using high K+ buffer with a negative correlation coefficient of -0.1699 to evoke Met-Enk release, suggesting that the limited release observed in **D** is due to modulation of peptide release rather than depletion of reserves. Data in **A-E** are transformed to a log scale and n=24 animals. In panel **A**, 2-way ANOVA analysis shows a main effect of peptide, treatment, and interaction. The *p*-values reported were calculated using a Sidak’s multiple comparisons test. **F.** Schematic describing the viral strategy and probe placement for the fiber photometry experiment in pEnk-Cre mice injected with the calcium sensor GCaMP6s in the NAcSh. **G.** Average z-score trace following the first, second, and third events involving experimenter handling. **H**. Heatmaps showing individual mouse z-scored fiber photometry responses before and after experimenter handling. **I**. Bar graphs showing the averaged z-score responses before experimenter handling and after experimenter handling (n=10 for **G-I**).

To test whether our findings regarding peptide release correlated with neuronal activity following the same stressor, we also monitored enkephalinergic neuron activity using fiber photometry following experimenter handling. To do this, we stereotaxically injected a Cre-dependent adeno-associated virus containing the calcium sensor GCaMP6s in the NAcSh of preproEnkephalin-Cre (pEnk-Cre) mice and implanted fiber photometry probes (**Fig. 2F**). During handling, we observed increased calcium activity in enkephalinergic neurons, even after it was repeated twice within a few minutes (**Fig. 2G**). Individual responses to experimenter handling varied appreciably, as shown by the heatmaps and bar plots (**Fig. 2H-I**); such variation could be due to slight differences in probe placement (**Suppl. Fig 3B,D**) as peptidergic neurons in the dNAcSh encode reward while neurons in the vNAcSh encode aversion [14,36]. Furthermore, the enkephalin precursor is expressed across the dorsal-ventral axis as shown in the *in situ* hybridization atlas (Allen Brain Institute). The fiber photometry data and peptide release data are concordant, as enkephalinergic neurons are activated following experimenter handling, which drives Met- and Leu-Enk somatodendritic release. However, the photometry and peptide release data differ in that repeated neuronal activation may not necessarily lead to repeated peptide release, as suggested by the decrease in peptide release over time in **Fig. 2D**. Additionally, fiber photometry cannot provide information about Met- and Leu-Enk independently, as it relies on the precursor gene, *penk*. Taken together, our data highlight the value and importance of coupling fiber photometry results with in vivo measurements of neuropeptides to determine release dynamics.

### Predator odor causes the release of Met-Enkephalin in the Nucleus Accumbens Shell

We wanted to test another stressor that is similar in nature to experimenter handling as the experimenter represents aversive visual, olfactory, and touch stimuli to the mouse. Fox odor has been widely demonstrated to be aversive to rodents and other mammalian species [37–39]. The benefit of using predator odor is that it represents what mice would encounter in the wild, outside of the lab environment. It also allows us to probe conserved circuits in laboratory mice related to aversion, threat, and defensive behaviors. Prior reports have shown that predator odor leads to an increase in the expression of the neuronal activation-associated transcription factor fos in enkephalin-positive neurons in different brain regions including the NAcSh [40,41]. Therefore, we sought to determine whether enkephalins are released in the NAcSh following exposure to fox urine. We first measured calcium activity of enkephalinergic neurons using fiber photometry after exposure to fox urine (**Fig. 3A**). We show that upon first exposure, the activity of enkephalinergic neurons is markedly increased (**Fig. 3B-D**). However, this enkephalinergic activity is significantly attenuated during second and third exposures that are separated by five minutes suggesting habituation to the stimulus (**Fig. 3B-D**). To determine whether the response to fox urine may also be due to the novelty of the weigh boat used to introduce the fox urine, we measured enkephalinergic calcium activity after introducing a weigh boat containing water to the home cage (**Suppl. Fig. 4A**). Interestingly, we show that enkephalinergic neurons also respond to the introduction of a novel object, and the response to the water-containing weigh boat is attenuated upon repeated exposure, which is what we observed for fox urine (**Suppl. Fig. 4B-D**). We compared the enkephalinergic neuron activation in response to both water and fox urine for each individual mouse that experienced both, and we see similar calcium transient responses to both the introduction of a novel object and the odor of a predator, suggesting the same subpopulation of enkephalinergic neurons may have been activated (**Suppl Fig. 4E**). Interestingly and importantly, only recently have researchers shown that a higher level of spiking in the striatum is not always reflected in greater photometry transients [42]. In the future, it will be important to determine whether the fox urine response indeed shows higher action potential spiking than the weigh boat with water. Although the calcium responses to both the fox-urine and water-containing weigh boats were similar in magnitude, it may be that different subpopulations of enkephalinergic neurons are activated in the NAcSh. Further exploration of the specific neurons that respond to the introduction of a novel object or the odor of a predator is warranted. We concluded that enkephalinergic neurons are activated by fox odor using fiber photometry. We were then interested in testing whether activation of enkephalinergic neurons leads to release of Met- and Leu-Enk, so we measured peptide levels in the NAcSh after fox odor using our method of microdialysis and nLC-MS (**Fig. 3E**).

**Figure 3.**
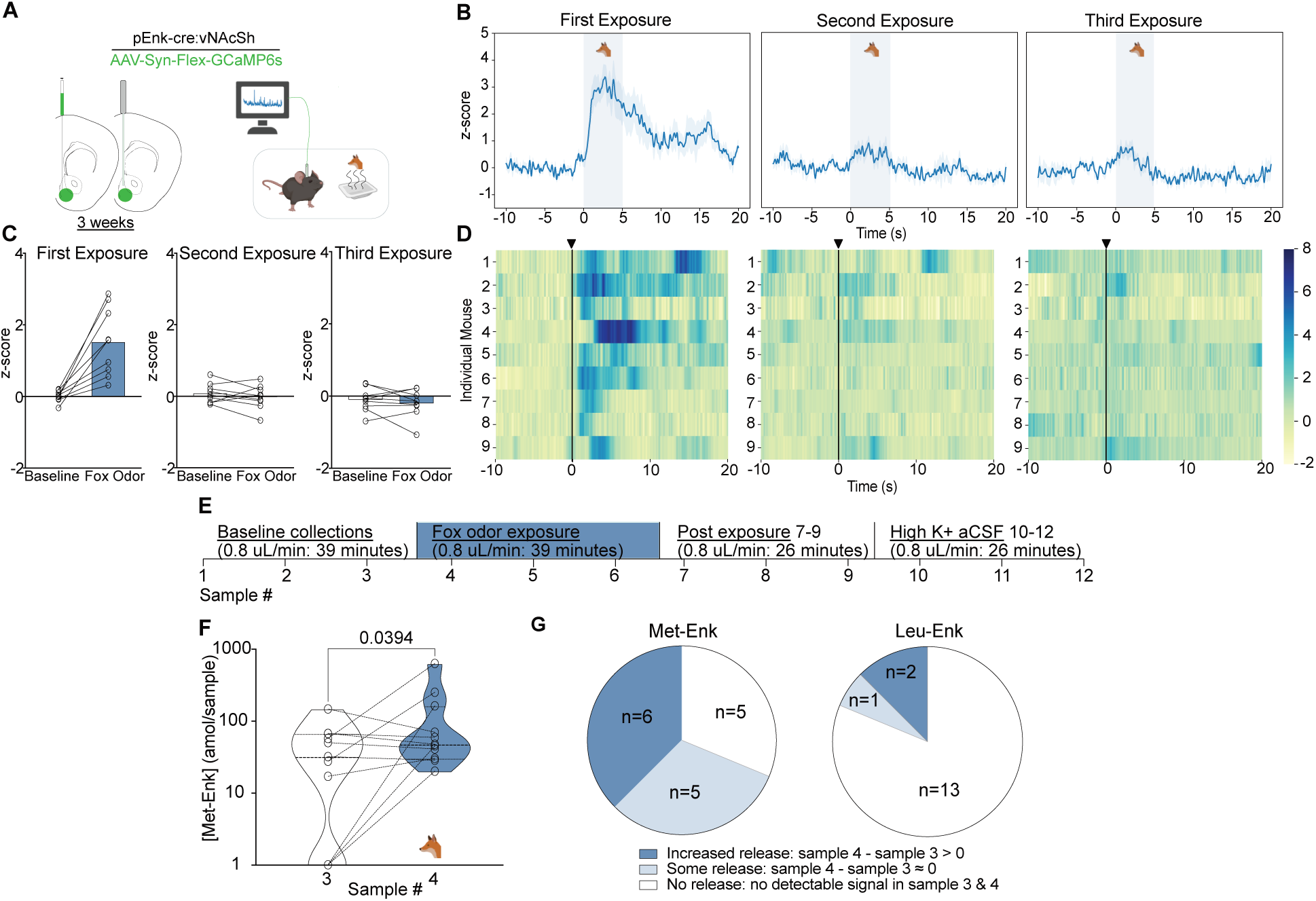
Fox odor exposure activates enkephalinergic neurons and drives the release of Met-Enk in the NAcSh. **A**. Viral strategy and probe placement for the fiber photometry experiment in pEnk-Cre mice injected with the calcium sensor GCaMP6s in the NAcSh. **B.** Averaged z-score traces of the first, second, and third exposure to fox odor. **C.** Bar graphs showing the averaged z-scored fiber photometry responses before and after exposure to fox odor **D**. Heatmaps showing individual mouse z-scored fiber photometry responses before and after exposure to fox odor. (n=9, for **A-D**) **E**. Experimental timeline describing the fox odor microdialysis experiment. **F**. There is a significant increase in Met-Enk concentration during fox odor exposure (sample 4), in comparison to sample, 3 before odor exposure. Data is transformed to a log scale and the *p*-value was calculated using a two-tailed paired t-test (n=11), t=2.369, df=10, mean difference 0.1388, 95% confidence interval -0.5419 to 0.2644 **G.** Pie charts showing the variation across animals in Met-Enk and Leu-Enk release profiles in response to exposure to fox odor, suggesting that such exposure may selectively cause the release of Met-Enk but not Leu-Enk. Increased release showed higher concentration of Met-Enk during exposure to fox odor (sample 4) than the sample before (sample 3), some release showed a comparable level to the sample before exposure, and no release showed no quantifiable concentration during exposure to fox odor.

In this separate group of animals, we show that there is a significant increase in Met-Enk release after exposure to fox odor (**Fig. 3F**). The response to predator odor varies appreciably: the responses are almost evenly split between elevated release, release with no elevation, or no detectable release (**Fig. 3G**). Enkephalins are thought to mediate resilience to stress, so the individual differences in responses to predator odor may indicate differences in stress adaptability [32,33,43,44]. Moreover, no sex differences in the responses to fox urine were observed, similar to responses to experimenter handling (**Suppl. Fig. 2B**). It is also worth noting that we only observed Leu-Enk release in 3 of 16 experiments. This is likely due to Leu-Enk not being released following predator odor or because it is released at a level below the LLOQ, as suggested by the observed 3:1 ratio of Met-Enk to Leu-Enk (**Fig. 3G**).

## Discussion

In summary, this work allows for multiple unique advances in studying endogenous opioids and neuropeptides in general. Although there has been substantial progress in fluorescent sensor-based techniques, electrochemical approaches, and immunoaffinity based assays, the investigation of opioid peptides has lagged due to their unique properties: sequence similarity, diversity, and low concentrations. Here, we address these challenges by advancing microdialysis coupled with nLC-MS. First, we have refined a method of methionine stabilization [21,45,46]that is transferrable to other neuropeptides and have decreased the LLOQ and temporal resolution, which are typically the main critiques of microdialysis and LC-MS approaches. As we have shown, if Met-Enk is not fully oxidized prior to LC-MS analysis, it is hard to quantify because of the presence of differentially oxidized species. The diversity of the signal detected impedes the analysis of a single sample and may lead to inaccurate concentration values. Here, we show that the oxidation of methionine allows for the stable detection of not only Met-Enk but also the simultaneous and selective detection of Leu-Enk.

Second, we quantified the relationship between Leu- and Met-Enk and offer a definitive *in vivo* ratio (3:1 Met-Enk to Leu-Enk) that has long been in question; this ratio corroborates human data [28] and suggests that mice are a useful model system for the study of enkephalinergic release dynamics. Based on this, future studies investigating the use of opioid peptides as biomarkers for psychiatric conditions will be readily translatable from rodents to humans and will allow for ease in reverse translation efforts. There is already some demonstrated interest in measuring opioid peptide levels in humans due to their roles in mood disorders, substance use, and pain [47,48]. Moreover, prior to our study, the predicted ratio of Met- to Leu-Enk was predicted based on the mRNA transcript sequence, so we are the first to confirm that post-translational cleavage indeed yields the predicted ratio. We also noted that Leu-Enk showed a greater fold increase relative to baseline after depolarization with high K+ buffer as compared to Met-Enk. This may be due to increased Leu-Enk packaging in dense core vesicles compared to Met-Enk or since there are two distinct precursor sources for Leu-Enk, namely both proenkephalin and prodynorphin while Met-Enk is mostly cleaved from proenkephalin (see Table 1) [49]. Third, we identify a role for the NAcSh in the acute stress response by demonstrating local release of Met- and Leu-Enk after stress exposure using our improved microdialysis and nLC-MS method. Although studies have suggested a role for the NAcSh in stress through measurements of *penk* transcripts, it had not been definitively shown that there is local release of enkephalins after stress exposure [35,50]. Our findings showed a robust increase in peptide release at the beginning of experiments, which we interpreted as due to experimenter handling stress that directly precedes microdialysis collections. However, there are other technical limitations to consider such as the fact that we were collecting samples from mice that were recently operated on. Another consideration is that the circulation of aCSF through the probe may cause a sudden shift in oncotic and hydrostatic forces, leading to increased peptide release to the extracellular space. As such, we wanted to examine our findings using a different technique, so we chose to record calcium activity from enkephalinergic neurons - the same cell population leading to peptide release. Using fiber photometry, we showed that enkephalinergic neurons are activated by stress exposure, both experimenter handling and fox odor, thereby adding more evidence to suggest that enkephalinergic neurons are activated by stress exposure which could explain the heightened peptide levels at the beginning of microdialysis experiments. Repeated experimenter handling led to repeated activation of enkephalinergic neurons. However, fox odor exposure showed significant attenuation after the first exposure. This is consistent with prior studies showing fast habituation to olfactory stimuli in rodents after initial exposure [37–39,51]. We also showed using fiber photometry that the novelty of the introduction of a foreign object to the cage, before adding fox odor, was sufficient to activate enkephalinergic neurons. This is consistent with the object being a stressor as it is newly introduced to the homecage. Alternatively, it could also mean that the stress response is more nuanced and is impacted by the novelty of the stressor. This is an area of interest to our group, and we will design future experiments to disentangle the novelty from the stressor in order to understand the enkephalinergic system better. Given that previous studies implicating enkephalins in stress used mRNA transcript measurements of proenkephalin, it was not possible to determine whether Leu- and/or Met-Enk were involved in the acute stress response. Our study suggests that both peptides are involved in experimenter handling stress, however this is less clear following exposure to fox odor where Met-Enk appears to be primarily involved. Most of our experiments with fox odor do not show Leu-Enk release, however, that could be due to Leu-Enk levels being below the LLOQ since we have demonstrated that it is 3 times lower than Met-Enk. Fourth, we show that Met- and Leu-Enk are released during acute stress in the NAcSh, offering evidence for their role in gating the stress response, which was suggested based on studies of enkephalins in the Locus Coeruleus [32]. Therefore, combined with other studies, our work suggests that the role of enkephalins in stress is generalizable to other brain regions, namely the ventral striatum. Based on our findings, it remains unclear how enkephalins modulate the stress response. It will be interesting to investigate whether enkephalinergic signaling modulates stress coping behaviors. Future studies will focus on determining the effect of the local release of Met- and Leu-Enk in the NAcSh on responses to stress in rodents.

Finally, when a limited amount of peptide is released over time, we show that peptide release is biologically constrained and can be experimentally driven using high K+ buffer or photo or chemogenetic stimulation [13]. We also show that data from fiber photometry corroborate peptide release data; however, fiber photometry does not offer the necessary specificity to correlate activation with release, or the distinction between different peptides that arise from the same precursor molecule. Additionally, it is important to stress that our fiber photometry studies used a calcium sensor as a proxy for neuronal activity and the temporal scale was on the order of seconds. A recent study using an opioid-peptide fluorescent sensor shows temporal scales on the order of minutes, which mirrors microdialysis timelines for neuropeptide detection, and offers evidence that neuropeptide dynamics are generally much slower than neurotransmitters *in vivo* [52]. In conclusion, this improved peptide detection method enables biological advances through an optimized approach for Met- and Leu-Enk detection *in vivo* with broad applicability to other neuropeptides.

## Materials and Methods

### List of Abbreviatons

FA: formic acid MeCN, acetonitrile
MeOH: methanol
DIA: Data-Independent Acquisition
>AGC: Automatic Gain Control
*nano*-LC-MS: capillary liquid chromatography interfaced to a mass spectrometer
MS1: mass spectra of peptide precursors
MS2: fragmentation mass spectrum of peptide from precursor ion

## Reagents

Acetonitrile (MeCN; J.T. Baker, cat. no. 9829-03)

Water (J.T. Baker, cat. no. 4218-03)

Formic Acid (FA; Sigma-Aldrich, cat. no. 56302)

Hydrogen Peroxide (Fisher Chemical, cat. no. H325-100)

Methanol (MeOH; Fluka, cat. no. 34966)

## Equipment

Stage tips

Laboratory pipetting needles with 90° blunt ends, 16-gauge, 2-inch length; Cadence Science, cat. no. 7938)

C18 extraction disks, diam. = 47 mm, 20 pack; (Empore, cat. no. 66883-U)

Adapter for stage tipping (Glygen, cat. no. CEN.24)

1.5-ml tubes (Axygen, Cat. No. MCT-175-C)

Eppendorf centrifuge (Eppendorf, model 5424R)

Vortex (Labnet VX100)

Autosampler vials (Sun-Sri, Cat. No. 200046)

Autosampler vial caps (Sun-Sri, cat. no. 501 382)

UPLC system for LC-MS of analysis of TMT-labeled peptides (EASY nL-LC 1000, Thermo Scientific)

Mass spectrometer for analysis of TMT-labeled peptides (Q-Exactive™ Plus Hybrid Quadrupole Orbitrap™, Thermo Scientific)

EASY-Spray™ column, 75 µm x 50 cm PepMap (Thermo Scientific™, cat. no. ES903)

## Animals

Adult male and female C57BL/6 mice at 8-15 weeks of age were used for microdialysis procedures. For fiber photometry, adult preproEnkephalin-Cre (pEnk-Cre) [53] male and female mice at ages between 15-24 weeks of age were used to measure enkephalinergic cell activity. Mice were group housed and kept at a 12-hour light/dark cycle with *ad libitum* access to water and food. The holding facility was both temperature and humidity controlled. All surgical and behavioral procedures were approved by the Washington University School of Medicine animal care guidelines in accordance with federal animal use regulations.

## Chemicals

Met and Leu Enkephalin standards including isotopically labeled forms were synthesized at New England Peptide and can be ordered by the research community. The heavy isotope labeled Leu Enkephalin is synthesized with a fully labeled L-Leucine (^13^C_6_, ^15^N) at the C terminus of the peptide (YGGF**L**). For Met Enkephalin, a fully labeled L-Phenylalanine (^13^C_9_, ^15^N) was added (YGG**F**M). The resulting mass shift between the endogenous (light) and heavy isotope-labeled peptide are 7Da and 10Da, respectively. Heavy isotope amino acids are guaranteed >99% pure. Peptide purity specifications are set at ≥ 95% pure by area on a regular HPLC gradient with UV detection. Artificial cerebrospinal fluid (aCSF) contained 124 mM NaCl, 2.5 mM KCl, 2 mM CaCl_2_, 1 mM MgCl_2_, 1.25 mM NaH_2_PO_4_, 24 mM NaHCO_3_, 5 mM HEPES, 12.5 mM Glucose adjusted to pH 7.4 with NaOH. High K^+^ ringer solution contained 48 mM NaCl, 100 mM KCl, 2.4 mM CaCl_2_, 0.85 mM MgCl_2_ adjusted to pH 7.4 with NaOH as described in (Al-Hasani, 2018). All salt compounds were purchased from Sigma (St. Louis, MO).

## *In vivo* microdialysis

Custom microdialysis probes (BASI Inc. part MD-2206) of 5 mm shaft length and 1 mm Polyacrylonitrile membrane with a 30 kDa molecular weight cutoff were used for the *in vivo* experiments. The probes were flushed prior to surgical implantation for 15 minutes at 10 μL/min with artificial cerebrospinal fluid as directed by the manufacturer using a BASI syringe pump (MD 1001). For the surgical procedure, mice were briefly anaesthetized in an induction chamber at 3% isofluorane before being placed in a stereotaxic frame (Kopf) secured with ear and bite bars. During the surgery, the isofluorane level was kept at 1.5-2%. The probe was then inserted into the ventral Nucleus Accumbens shell (stereotaxic coordinates from bregma: +1.3 [AP], ±0.5 [ML], −5.0 mm [DV]). Implanted probes were secured with dental adhesive (C&B metabond kit) followed by cyanoacrylate (Loctite superglue gel control).

After the completion of the surgery, each mouse was allowed to recover for 30 minutes. The mouse was then connected to an inlet perfusion line to circulate aCSF or high K+ ringer solution and an outlet perfusion line to collect the interstitial fluid from the NAcSh. The outlet line directly collected sample in a tube that is kept at 4 °C throughout the experiment. The flow of aCSF was then then turned on and sample collection time started as sample began to collect in the outlet tube. The flow rate of aCSF was set to 0.8 μL/min during sample collection. Fractions were collected every 13 minutes resulting in a 10 μL sample. After each collection, the samples were spiked with 2 μL of 12.5 fmol isotopically labeled Met-Enkephalin and Leu-Enkephalin. The microdialysis rig (BASI Raturn MD 1404) was equipped with bedding, regular mouse chow and water ad libitum. While connected to the perfusion lines, the mice were allowed to move freely. For the experiments measuring evoked release, the high K+ ringer solution was perfused through the lines instead of aCSF at 0.8 μL/min and fractions were collected every 13 minutes as described previously. The samples were similarly spiked with 2 μL of 12.5 fmol isotopically labeled Met-Enkephalin and Leu-Enkephalin.

For the experiments where the animals were exposed to the fox urine odor, 3 mL of fox urine was absorbed into a kim wipe contained in a chemical weigh boat. The weigh boats were prepared and remained in a fume hood the experiment to prevent the odor permeating the experimental space before the planned collection. Then, the experimenter placed the weigh boat in the microdialysis rig and allowed the mouse to explore it. The exposure was standardized to span 3 fractions (39 minutes) during each of the experiments while aCSF was being perfused through the lines.

## In vivo sample processing

### Methionine Oxidation Reaction

For met-enkephalin detection and stabilization, the animal samples underwent a methionine oxidation reaction [21]. To each sample vial, 28 μl of 0.1% (vol/vol) formic acid (FA) was added and then the samples were transferred to 0.5 mL tubes from the collection tubes. Afterwards, 40 μl of a mixture of 6% FA (vol/vol) and 6% hydrogen peroxide (vol/vol) was added to each of the sample tubes and mixed and gently spun. The samples were then incubated overnight at room temperature.

### Solid Phase Extraction with C18 stage tips

After the methionine oxidation reaction, a solid phase extraction process was performed using C18 stage tips. Stage tips were conditioned before sample loading. To each of the stage tips, 100 μl of methanol is added and then spun for 2 minutes at 2,000 rpm in microcentrifuge tubes. The stage tips were then rinsed with 100 μl of a mixture of 50% acetonitrile (MeCN; (vol/vol)) and 0.1% FA (vol/vol) followed by a 2-minute spin at 2,000 rpm. This was followed by a rinse in 0.1% FA (vol/vol) and a 2-minute spin at 3,000 rpm. The final rinse was done by adding 100 μl of 1% FA (vol/vol) to each of the stage tips. Following the final rinse, the samples were added to the C18 stage tips and spun for 10 minutes at 0.8 rpm. After ensuring that the samples passed through the stage tips, the samples were desalted with 0.1% FA (vol/vol) and spun for 1.5 minutes at 3,000 rpm twice. After the spins were done, the stage tips were directly placed into autosampler vials to elute the analyte with 60 μl of 50% MeCN, 0.1% FA (vol/vol) using a syringe. Eluates in the autosampler vial were dried in a speedvac, reconstituted in 3 μl 1% FA (vol/vol) and set up on the autosampler for injection on the mass spectrometer.

### *Nano*-LC-MS/MS

The samples in formic acid (1%) were loaded (2.5 µl) onto a 75 µm i.d. × 50 cm Acclaim^®^ PepMap 100 C18 RSLC column (Thermo-Fisher Scientific) on an EASY *nano*LC (Thermo Fisher Scientific) at a constant pressure of 700 bar at 100% A (1%FA). Prior to sample loading the column was equilibrated to 100%A for a total of 11 µL at 700 bar pressure. Peptide chromatography was initiated with mobile phase A (1% FA) containing 5%B (100%MeCN, 1%FA) for 1 min, then increased to 20% B over 19 min, to 32% B over 10 min, to 95% B over 1 min and held at 95% B for 19 min, with a flow rate of 250 nL/min. The data was acquired in data-independent acquisition (DIA) mode. The full-scan mass spectra were acquired with the Orbitrap mass analyzer with a scan range of *m/z* = 150 to 1500 and a mass resolving power set to 70,000. Data-independent (DIA) high-energy collisional dissociations were performed according to the inclusion list with a mass resolving power set to 17,500, an isolation width of 2 Da, and a normalized collision energy setting of 27. The maximum injection time was 60 ms for parent-ion analysis and product-ion analysis. The automatic gain control (AGC) was set at a target value of 3e6 ions for full MS scans and 2e5 ions for MS2. The inclusion list contained 4 peptide entries. The loop count was set at 30. Retention time windows were set at 20 min. Each sample was run in two technical replicates and the peak area ratio was averaged before concentration calculations of the peptides were conducted. Several quality control steps were conducted prior to running the *in vivo* samples. 1) Two technical replicates of a known concentration were injected and analyzed – an example table from 4 random experiments included in this manuscript is shown below. 2) The buffers used on the day of the experiment (aCSF and high K+ buffer) were also tested for any contaminating Met-Enk or Leu-Enk signals by injecting two technical replicates for each buffer. Once these two criteria were met, the experiment was analyzed through the system. If either step failed, which happened a few times, the samples were frozen and the machines were cleaned and restarted until the quality control measures were met.

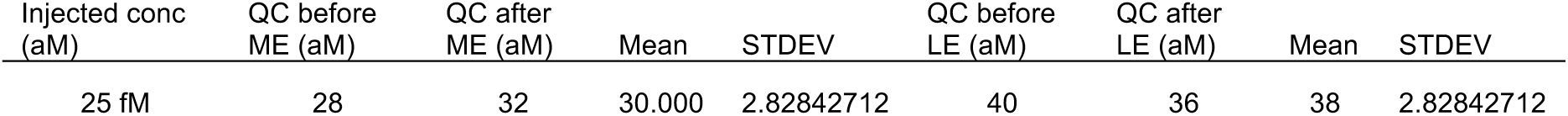
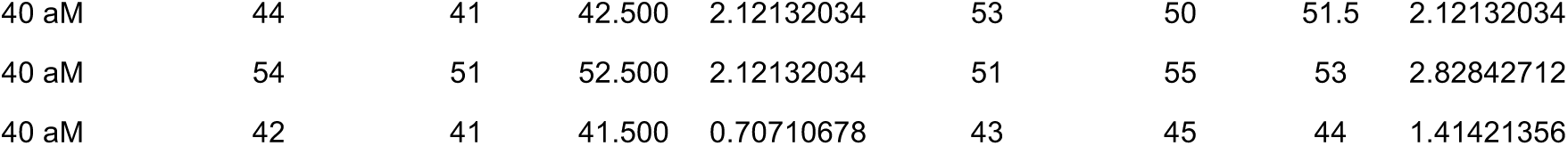

### Stable-isotope dilution assay for measurement of enkephalins

Peptides were synthesized as high purity natural abundance and stable isotope-labeled pairs for the development of a targeted proteomics method for enkephalins. Assays were developed as Tier 2 assays according to CPTAC guidelines [22]. Co-eluting peak boundaries for the internal standard and endogenous peptides were manually refined using the Skyline software package [54]. Integrated peak areas for the three most intense fragment ions were then exported for determination of peak area ratios of endogenous to internal standard signal, which were used as a basis for determining abundance and quantifying changes in enkephalin levels in mouse studies.

### *In vivo* fiber photometry

#### Stereotaxic surgery

For the surgical procedure, pEnk-Cre mice were briefly anaesthetized in an induction chamber at 3% isofluorane before being placed in a stereotaxic frame (Kopf) secured with ear and bite bars. During the surgery, the isofluorane level was kept at 1.5-2%. Then we performed a craniotomy and unilaterally injected using a Nanoject III (Drummond) 200-250 nL of AAV5.Syn.Flex.GCaMP6s.WPRE.SV40 (Addgene: 100845-AAV5) at the following coordinates :+1.3 [AP],±0.5 [ML], −5.0 mm [DV]. To allow for viral expression, the fiber photometry probes (5 mm shaft length, doric lenses) were implanted 3 weeks later at +1.3 [AP], ±0.5 [ML], −4.75 mm [DV]. Implanted probes were secured with dental adhesive (C&B metabond kit) followed by cyanoacrylate (Loctite superglue gel control).

#### Fiber photometry recording procedure

Before recording, the implanted probes were tethered to an optic fiber using a ferrule sleeve (Doric, catalog no. ZR_2.5). Two light-emitting diodes (LEDs) were used to excite GCaMP6s. A 331 Hz sinusoidal LED light (Doric) of wavelength 470 nm 470 ± 20 nm was used for the Ca^2+^ - dependent signal. A 210 Hz sinusoidal LED light of wavelength 405 ± 10 nm (Doric, LED driver) was used to evoke the Ca^2+^ -independent signal and serve as the isosbestic control. GCaMP6s fluorescence traveled through the same optic fiber before being bandpass filtered (525 ± 25 nm, Doric, catalog no. FMC4), transduced by a femtowatt photoreceiver (New Focus Model 2151) and recorded by a real-time processor (TDT, RZ5P). The signals were extracted by the Synapse software (TDT). During the recording, two behavioral manipulations were introduced. The first was after a 10-minute recording in a home cage environment, an experimenter handled the mouse while the recording was ongoing. The handling bout which mimicked traditional scruffing lasted about 3-5 seconds. The mouse was then let go and the handling was repeated another two times in a single session with a minimum of 1-2 minutes between handling bouts. Mice were habituated to this manipulation by being attached to the fiber photometry rig, for 3-5 consecutive days prior to the experimental recording. Additionally, the same maneuver was employed when attaching/detaching the fiber photometry cord, so the mice were subjected to the same process several times. The second behavioral manipulation was introduced to the same mice a week after the first recording session. After a 10-minute recording in the home-cage environment, a weighboat containing a kim wipe that had absorbed 3 mL of fox urine was placed in the cage as the recording proceeded. The weigh boat was removed after a minute of exploration then re-introduced into the cage for another two times, separated by 5 minutes from the last exposure, during the session. In a subset of the animals (5 of 9), the fox urine exposure was preceded by water exposure following the same process described above in the same recording session.

#### Fiber Photometry Data Analysis

Data analysis was performed through the python toolbox GuPPy [55]. Briefly, the isosbestic control signal was used to correct for motion and photobleaching artifacts. The control channel is fitted to the signal channel using a least squares polynomial fit of degree 1. Transient detection occurred after using a 15s moving window for thresholding transients. Then, high-amplitude events were identified and their timestamp of occurrence during the session was reported. To determine the effect of handling or fox urine exposure on enkephalinergic cell activity, the event timestamp was used to observe changes in activity before and after the behavioral manipulation. The average of all trials is then calculated for repeated exposures and for exposures in multiple animals. The fiber photometry data was represented as z-scores.

#### Immunohistochemistry and verification of probe placement

After the completion of the experiment, mice were anaesthetized with a mixture of ketamine, xylazine and acepromazine and transcardially perfused with ice-cold 4% paraformaldehyde in phosphate buffer. After the perfusion, brains were dissected and fixed for 24 hours in 4% paraformaldehyde at 4°C and then placed in 30% sucrose solution for cryoprotection for a minimum of 48 hours. The brains were then sliced into 30 um sections, washed three times in PBS and blocked in PBS containing 0.5% Triton X-100 and 5% goat serum for one hour. Following the blocking step, the slices were stained for the glial fibrillary acidic protein (GFAP), an astrocyte marker to determine glial scarring around the probe site (guinea pig anti-GFAP, synaptic systems) at 1:500 dilution and incubated at 4°C for 16 hours. The next day, the sections were washed three times with PBS and stained with the secondary antibody, mouse anti-guinea pig, Alexa Fluor 594 (Invitrogen) at 1:500 dilution and incubated for two hours at room temperature. After the incubation, the slices were washed three times in PBS, followed by two washes with PB. Finally, the slices were mounted on glass slides with Hard set Vectashield with DAPI staining (Vector Labs) and imaged using an epifluorescence microscope (Leica DM6 B). Correct probe placement in the NAcSh is represented in **supplementary figure 1B**. For the verification of the injection and probe placement for fiber photometry, the same procedure described above was performed for tissue preparation. After slicing, the slices were washed in a 0.1% Triton for one hour in order to permeabilize the cells followed by two 10-minute PBS washes. The slices were then stained with Neurotrace (Nissl) red (Invitrogen) to mark neurons by adding 5 μL to each well for 2 hours at room temperature followed by 3 10-minute PBS washes and 3 10-minute PB washes. Finally, the slices were mounted using Hard set Vectashield (Vector Labs). GCaMP6s expression was determined by observing green fluorescent protein expression colocalized with Neurotrace staining. Correct probe placement in the NAcSh is represented in **supplementary figure 1D**.

### Statistical Analysis

We performed simple linear regression analysis on the forward and reverse curves and reported the line equations. For the forward curves, the regression was applied to the measured concentration of the light standard as the theoretical concentration was increased. For plotting purposes, we show the measured peak area ratios for the light standards in the forward curves. For the reverse curves, the regression was applied to the measured concentration of the heavy standard, as the theoretical concentration was varied. The proteomics data is represented as violin plots with individual points depicted and dashed lines indicating quartiles. The middle-dashed lines represent the median. The violin plots were created using Graphpad Prism. The data was log_10_ transformed to reduce the skewness of the dataset caused by the variable range of concentrations measured across experiments/animals. Prior to log transformation, the measurements failed normality testing for a Gaussian distribution. After the log transformation, the data passed normality testing, which provided the rationale for the use of statistical analyses that assume normality. Two-way ANOVA testing with peptide (Met-Enk or Leu-Enk) and treatment (buffer or stress for example) as the two independent variables. Post-hoc testing was done using Šídák’s multiple comparisons test and the *p* values for each of these analyses are shown in the figures (Figs. 1F, 2A). A paired t-test was performed on the predator odor proteomic data before and after odor exposure to test that hypothesis that Met-Enk increases following exposure to predator odor (Fig. 3F). These analyses were conducted using Graphpad Prism. To determine the linear relationship between the levels of Met-Enk to Leu-Enk in the same samples, we conducted simple linear regression analysis and reported the slopes of the lines. This is based on prior literature positing a relatively fixed ratio of Met-Enk to Leu-Enk, and we corroborate that here [25–27]. For the fiber photometry data, the z-scores were calculated as described in using GuPPy which is an open-source python toolbox for fiber photometry analysis [55]. The z-score equation used in GuPPy is z=(ΔF/F-(mean of ΔF/F)/standard deviation of ΔF/F) where F refers to fluorescence of the GCaMP6s signal. For the averaged plots depicted in Fig. 2 and Fig. 3, z-scores from all individual animals were averaged and shown as one trace with the SEM as the highlighted area. The purpose of the averaged traces is to show the extent of concordance of the response to experimenter handling and predator odor stress among animals with the SEM demonstrating that variability. The heatmaps depict the individual responses of each animal. The heatmaps were plotted using Seaborn in Python and mean traces were plotted using Matplotlib in Python.

## Author Contributions

- Conceptualization: RA, MOM, RRT, RS
- Methodology: MOM, RA, PEG, RC, JM, RS
- Investigation: MOM, PEG, RC, SMC, JW
- Formal Analysis: MOM, PEG, RS
- Visualization: MOM, JM
- Supervision: RA, RRT, RS
- Writing—original draft: MOM, PEG, RS
- Writing—review & editing: RA, MOM, PEG, RC, SMC, JM, JW, RS, RRT

## Acknowledgments

The figure schematics were created with biorender.com. We would like to thank the Moron-Concepcion lab for providing us with the Enk-Cre mouse line. We would like to thank Justin W. Baldwin for his guidance on appropriate data visualization tools and for his support in proofreading the manuscript. We would also like to thank Dr. Jordan McCall for his advice on data analysis and visualization as well as proofreading the manuscript. Thank you to Emily Wheeler for proofreading the manuscript and providing feedback on sentence structure and flow. We would also like to thank the entirety of the proteomics core team including Dr. Yiling Mi for her assistance. Thank you to the entirety of the Al-Hasani and McCall labs for their feedback on experimental design and analysis. Finally, thank you to Dr. Tim Holy, Dr. Robyn Klein, and Dr. Alexxai Kravitz for their feedback.

