## Supplemental Information for "Highly sensitive *in vivo* detection of dynamic changes in enkephalins following acute stress"

### Supplementary Materials

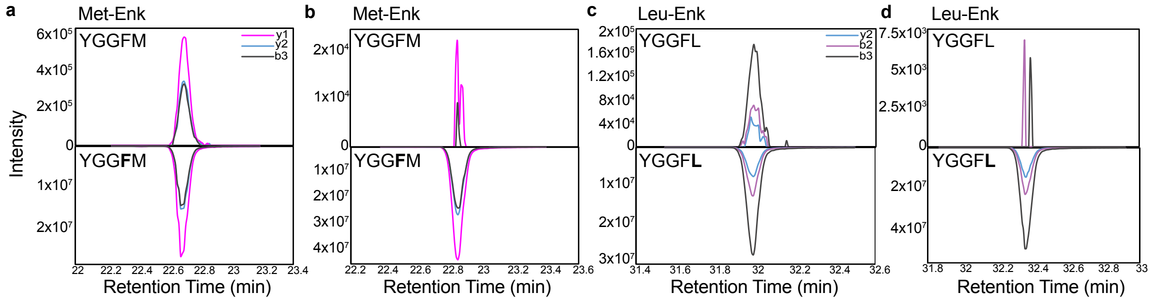


**Supplementary figure 1**: **Technical advancements enabled using internal standards for Met- and Leu-Enk detection. a**. A representative trace of high signal-to-noise endogenous Met-Enk signal (top) compared to the isotopically labeled internal standard (bottom). A representative low signal-to-noise trace of endogenous Met-Enk signal (top) compared to the isotopically labeled standard (bottom). **c.** A representative high signal-to-noise trace of endogenous Leu-Enk signal (top) compared to the isotopically labeled internal standard (bottom). **d.** A representative low signal-to-noise trace of endogenous Leu-Enk signal (top) compared to the isotopically labeled standard (bottom). The bolded amino acids in the sequences for Met- and Leu-Enk represent the isotopically labeled standards.

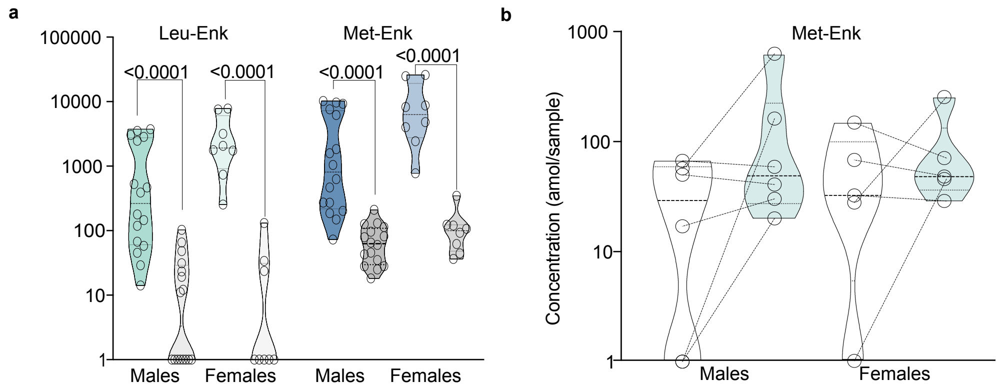


**Supplementary figure 2**: **There are no significant differences between male and female responses to handling or fox odor exposure**. **a**. Male and female responses to handling are comparable for both Leu- and Met-Enk. (n=16 males, n=8 females). The 3-way ANOVA analysis showed a significant main effect of sex ($p$= 0.0049) and sex*treatment interaction ($p$= 0.0107). However, after correcting for multiple comparisons, there were no differences between male and female conditions. **b.** Met-Enk release is comparable between males and females as the main of effect of sex in the 2-way ANOVA is not signifcant ($p$= 0.6821) and neither is the sex*treatment interaction ($p$= 0.5504). (n=6 males, n=5 females).

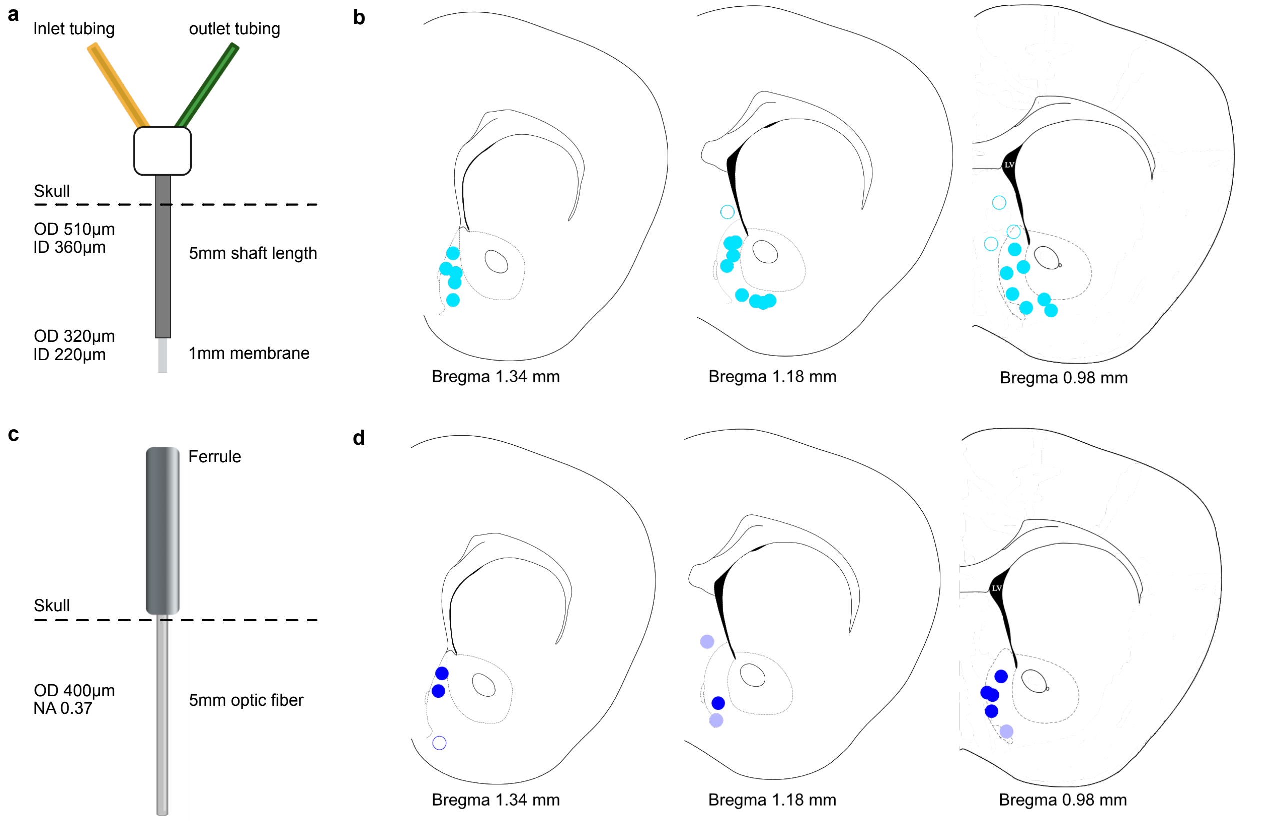


**Supplementary Figure 3**: **Hit map of microdialysis and fiber photometry probes**. **a.** Microdialysis probe dimensions including membrane size and inner and outer diameters (ID and OD respectively). **b.** Hit map of microdialysis probes represented in main figures. Full circles represent hits and empty circles represent misses. **c.** Dimensions for fiber photometry probes including OD of optic fiber and numerical aperture (NA). **d.** Hit map of placement of fiber photometry probes. Dark circles represent hits, light circles represent near hits when activity was still recorded, empty circles represent misses.

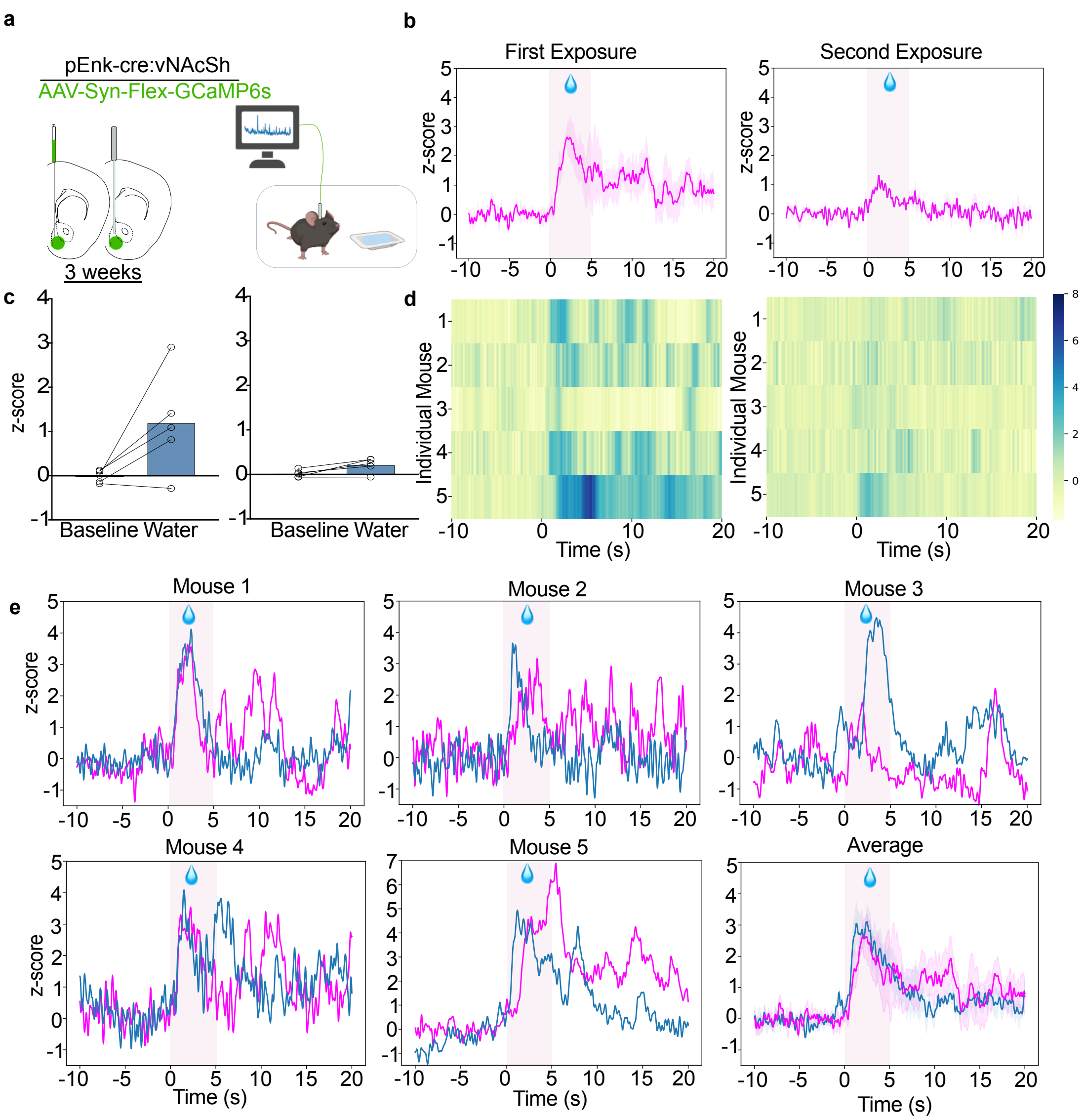


**Supplementary figure 4: Water exposure activates enkephalinergic neurons in the NAcSh. a**. Viral strategy and probe placement for the fiber photometry experiment in pEnk-Cre mice injected with the calcium sensor GCaMP6s in the NAcSh. **b.** Averaged z-score traces of the first and second water exposures. The shaded region represents the sem. **c.** Bar graphs showing the averaged z-scored fiber photometry responses before and after exposure to water. **d**. Heatmaps showing individual mouse z-scored fiber photometry responses before and after water exposure. (n=5, for **a-d**) **e**. Individual mouse responses to fox urine (blue) and water exposure (magenta) plotted on the same graph for comparison. The final graph is the averaged data for all 5 mice with the shaded region representing SEM.
